# Repurposing the yeast peroxisome to compartmentalize a toxic enzyme enables improved (*S*)-reticuline production

**DOI:** 10.1101/2020.03.23.000851

**Authors:** Parbir S. Grewal, Jennifer A. Samson, Jordan J. Baker, Brian Choi, John E. Dueber

**Affiliations:** Department of Chemical & Biomolecular Engineering, University of California, Berkeley, CA 94720, USA; Department of Bioengineering, University of California, Berkeley, CA 94720, USA; UC Berkeley and UCSF Graduate Program in Bioengineering, University of California, Berkeley, CA 94720, USA; Biological Systems & Engineering Division, Lawrence Berkeley National Laboratory, Berkeley, California, USA

## Abstract

Eukaryotic cells compartmentalize metabolic pathways in organelles to achieve optimal reaction conditions and avoid crosstalk with other factors in the cytosol. Increasingly, engineers are researching ways in which synthetic compartmentalization could be used to address challenges in metabolic engineering. Here, we identified that norcoclaurine synthase (NCS), the enzyme which catalyzes the first committed reaction in benzylisoquinoline alkaloid (BIA) biosynthesis, is toxic when expressed cytosolically in *Saccharomyces cerevisiae* and, consequently, restricts (*S*)-reticuline production. We developed a compartmentalization strategy that alleviates NCS toxicity while promoting increased (*S*)-reticuline titer, achieved through efficient targeting of toxic NCS to the peroxisome while, crucially, taking advantage of the free flow of metabolite substrates and product across the peroxisome membrane. We identified that peroxisome protein capacity in *S. cerevisiae* becomes a limiting factor for further improvement of BIA production and demonstrate that expression of engineered transcription factors can mimic the oleate response for larger peroxisomes, further increasing BIA titer without the requirement for peroxisome induction with fatty acids. This work specifically addresses the challenges associated with toxic NCS expression and, more broadly, highlights the potential for engineering organelles with desired characteristics for metabolic engineering.

## Introduction

Cell factories, using production hosts ranging from bacterial, fungal, mammalian, and plant cells, are used to make important industrial products including pharmaceuticals, proteins, commodity chemicals, and biofuels^1–4^. As cellular engineering becomes increasingly sophisticated, incorporating longer and more complex heterologous metabolic pathways, the likelihood of crosstalk with native cellular functions in the host cell also increases. Consequently, there is greater risk for cytotoxicity and effects on host cell growth. Reduced growth has many negative consequences, including reduction in product titer due to fewer cells, decrease in production rate due to slower growth, and decrease in strain robustness because mutations that relieve toxicity are often ones that reduce or eliminate pathway flux.

Subcellular compartmentalization is an evolved strategy for coping with unproductive or harmful crosstalk^5–8^. In eukaryotes, lipid bilayer membrane-enclosed organelles are used to direct enzymatic activity toward specific substrates. In many cases, these activities would be unproductive in the cytosol. For example, vacuolar proteases have broad substrate specificity but their degradative activity is directed toward specific substrates due to their compartmentalization in the vacuole^9^. These degradation mechanisms are extremely precise. For instance, the Cvt pathway is known to degrade only three proteins, utilizing Atg19p to facilitate selective autophagy into the vacuole^10^. Additionally, many vacuolar proteases are imported as inactive forms called zymogens that are further processed once inside the vacuole into active forms, protecting proteins in the secretory pathway from degradation^11^.

Compartments are also used to sequester toxic metabolites. For example, the parasite *Trypanosoma brucei* respecializes the peroxisome during the part of its life cycle in the human bloodstream from the β-oxidation of long-chain fatty acids to organelles known as glycosomes where the early steps of glycolysis are compartmentalized. While most organisms use allosteric feedback regulation of these glycolytic enzymes to prevent toxic accumulation of intermediates, *T. brucei* instead uses a compartmentalization strategy to enable increased flux while preventing toxic intermediate accumulation in the cytosol^12,13^. In the context of biomanufacturing and synthetic biology, compartmentalization may prove to be a powerful strategy to isolate toxic heterologous components away from the host cell cytosol, thereby limiting cytotoxicity, improving growth, and increasing product titer.

Engineered compartmentalization has garnered considerable attention in recent years^7,14–18^. Many of the reports of engineered compartmentalization take advantage of the endogenous cofactor supply or substrate pool in a particular organelle. For example, the mitochondria has been used for its supplies of acetyl-CoA^19,20^ and α-ketoisovalerate^21^, the peroxisome for acyl-CoA^22,23^, farnesyl diphosphate^24^, and acetyl-CoA^18^, and the vacuole for halide ions and *S*-adenosylmethionine^25^. Organelles have also been harnessed for the reactions that they natively perform. For example, the endoplasmic reticulum and Golgi in Chinese Hamster Ovary cells are used for the glycosylation of heterologous proteins^26^, and recently the methanol assimilation pathway in the peroxisomes of *Pichia pastoris* was engineered into a carbon dioxide fixation pathway where six native peroxisome enzymes were utilized^17^. Advances in the development of synthetic compartments have also been reported, including the synthesis of prokaryotic encapsulin nanocompartments in eukaryotes^15,27^, bacterially-derived proteinaceous gas vesicles in mammalian cells^28^, and light-inducible phase-separated organelles in yeast^16^.

In this work, we discovered that a titer-limiting enzyme in the industrially-valued benzylisoquinoline alkaloid (BIA) family of plant natural products is toxic when expressed in a yeast production host. The BIA family consists of ∼2500 natural products exhibiting a variety of potential therapeutic bioactivities including antibacterial (berberine, sanguinarine), anticancer (noscapine), antispasmodic (papaverine), and analgesic (codeine, morphine)^29,30^. The first committed BIA molecule, (*S*)-norcoclaurine, is produced through the condensation of two tyrosine-derived substrates, dopamine and 4-hydroxyphenylacetaldehyde, by the enzyme norcoclaurine synthase (NCS)^31–33^. NCS is thought to be a gatekeeper enzyme that restricts entry into the BIA pathway^34,35^. Accordingly, high NCS activity is required for high production levels of BIAs in heterologous hosts. However, we found that our most active variant of NCS is toxic when expressed in the cytosol of *S. cerevisiae* and determined that the toxicity is caused by the NCS protein itself rather than by its catalytic function in the condensation of dopamine and 4-hydroxyphenylacetaldehyde to produce (*S*)-norcoclaurine.

We propose repurposing of the yeast peroxisome as the site for subcellular compartmentalization of toxic NCS enzyme. The peroxisome has several attributes that make it an ideal compartment for alleviating NCS toxicity without inhibiting flux through the BIA pathway. First, because peroxisomes in *S. cerevisiae* are used for the β-oxidation of fatty acids, they are not essential under typical industrial fermentation conditions where simple sugars are used as the carbon source^36^. Therefore, *S. cerevisiae* peroxisomes can be repurposed for use in applications orthogonal to their evolved function without having major effects on native cellular processes and culture viability^18,37^. Second, heterologous protein can be targeted to the peroxisome by simply appending a small peptide tag to the C-terminus (peroxisomal targeting signal type 1, PTS1)^38,39^ or N-terminus (PTS2)^40,41^ of the protein of interest. In fact, our lab previously developed an enhanced targeting tag (ePTS1) that considerably increased the rate of protein import into the peroxisome^37^. Third, the peroxisome is thought to contain the highest concentration of protein in the eukaryotic cell^42,43^ and can compartmentalize an impressive amount of heterologous cargo^37^, more than has been demonstrated for other organelles^44^. Fourth, peroxisomes are permeable to many small molecules under approximately 500-700 daltons, meaning that peroxisomally compartmentalized NCS would likely retain access to its substrates^37,45^.

Here, we demonstrate that compartmentalization of toxic NCS in the peroxisome can both alleviate cytotoxicity and improve product titer. We utilize this compartment to selectively partition the toxic protein from other cytosolic factors while taking advantage of the ability of the peroxisome to allow small molecule substrates and products to freely diffuse into and out of the peroxisome lumen. Compartmentalization of toxic NCS enzyme improves production of (*S*)-norcoclaurine and the key branchpoint intermediate (*S*)-reticuline. More broadly, we provide a demonstration of the utility of the peroxisome as a two-way insulating compartment, one that can protect the host cell from toxicity associated with heterologous protein expression and, conversely, can protect heterologous protein from native cellular processes such as protein degradation. Although the peroxisome naturally has an impressive cargo capacity compared to other organelles, improved capacity is still desired for many engineering applications, including the compartmentalization of a toxic, activity-limiting metabolic enzyme as described here. We demonstrate initial progress toward this goal by using engineered transcription factors to mimic the peroxisome proliferation effect that is produced by long-chain fatty acid induction and utilize this strategy to further improve BIA titer.

## Results

### Expression of most active NCS variant causes cytotoxicity in yeast

While working to connect yeast central metabolism to the heterologous benzylisoquinoline alkaloid (BIA) pathway of plant-derived drug molecules, our group previously identified the norcoclaurine synthase (NCS) enzyme as a key rate-limiting step preventing high titer production of BIAs (Fig. 1a)^46^. Since then, we have tested several homologues (and truncated variants of these homologues for optimized expression) to identify enzyme sequences capable of producing higher levels of the NCS product, (*S*)-norcoclaurine, when expressed in yeast. In comparison to our original NCS derived from *Papaver somniferum*, heterologous expression of the NCS homologue from *Coptis japonica* resulted in a 3-fold higher (*S*)-norcoclaurine titer (Supplementary Fig. 1). Moreover, truncation of the N-terminus of the *Coptis japonica* homologue to remove residues upstream of the Bet v1 domain, the critical fold of NCS proteins^33^, resulted in 20-fold higher titer than our original *Papaver somniferum* variant - the highest activity yet described in *S. cerevisiae*. For simplicity, we will refer to this truncated *Coptis japonica* homologue as tNCS. Previous studies have also reported improved titers upon truncation of NCS enzymes, with evidence suggesting that this effect is due to improved expression and/or stability of the protein^47–49^. (*S*)-Norcoclaurine titer was directly dependent on the expression level of tNCS, with stronger promoters and copy numbers resulting in higher (*S*)-norcoclaurine titers (Fig. 1b). Copy number was increased compared to chromosomal integration by using a CEN6/ARS4 yeast replicating plasmid (∼2-5 copies per cell^50^). Additionally, transcription levels were controlled via a range of previously characterized yeast promoters^51,52^. Unexpectedly, we also observed that heterologous expression of tNCS resulted in a reduction in yeast growth rate in an expression-level dependent manner (Fig. 1c). Moderate expression of tNCS (pRPL18B) had almost no effect on cell growth when compared to cells that do not express tNCS, whereas strong expression (pTDH3) considerably decreased both the growth rate and final cell density of the yeast culture.

**Figure 1.**
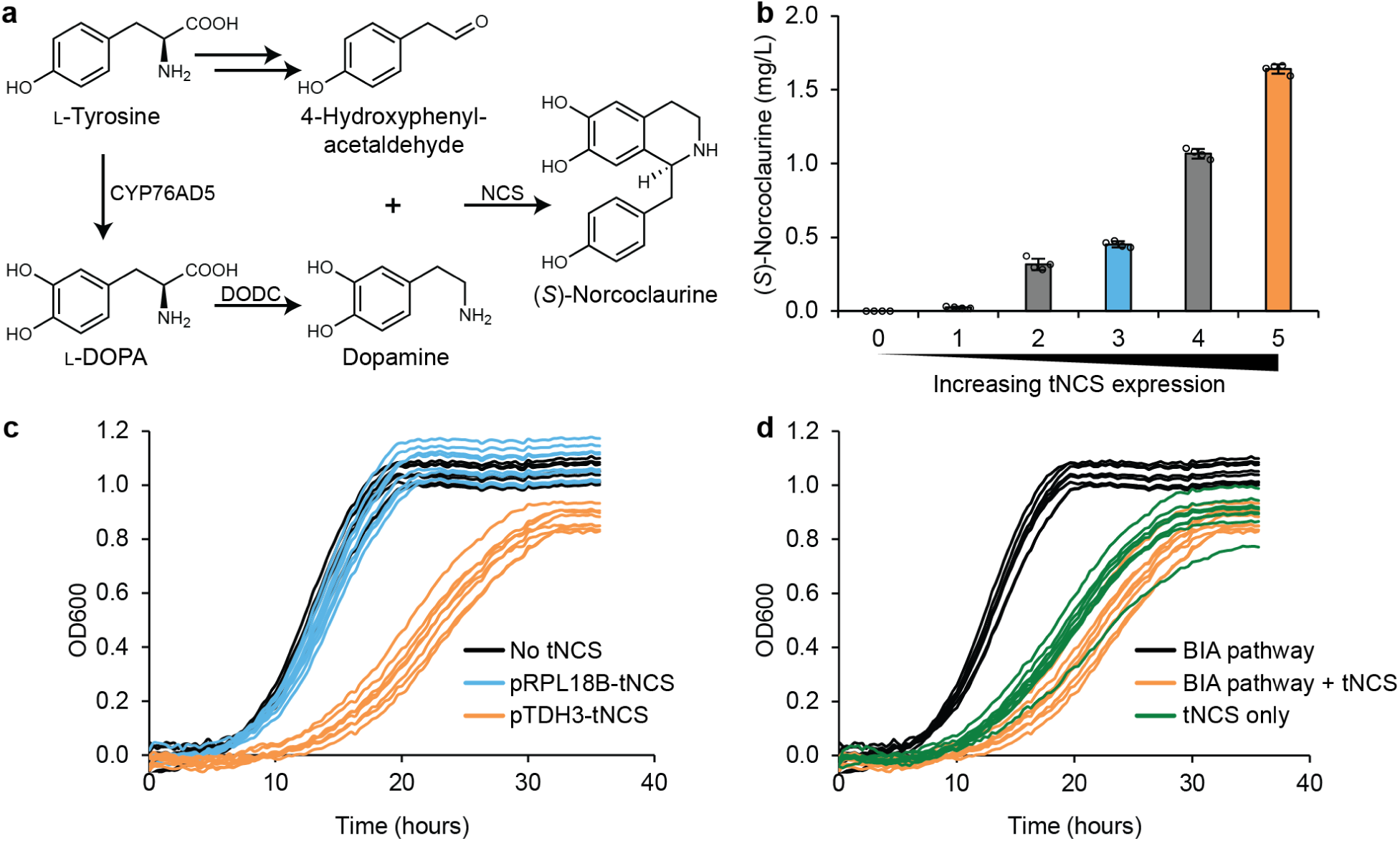
Cytosolic expression of truncated norcoclaurine synthase (tNCS) is toxic to *S. cerevisiae*. (**a**) Metabolic pathway for production of (*S*)-norcoclaurine from L-tyrosine using three heterologous enzymes: tyrosine hydroxylase (CYP76AD5), dopamine decarboxylase (DODC), and tNCS. 4-Hydroxyphenylacetaldehyde is produced endogenously by *S. cerevisiae*. (**b**) Higher expression of tNCS results in higher titer of (*S*)-norcoclaurine. The following well-characterized promoters were used to drive expression of tNCS on a CEN6/ARS4 plasmid: 1=pREV, 2=pRNR2, 3=pRPL18B, 4=pTEF1, 5=pTDH3. 0=No NCS. Error bars represent mean ± s.d. of four biological replicates. (**c**) Higher expression of tNCS results in slower growth, with significant toxicity observed at pTDH3 expression level. n=8 biological replicates per strain. (**d**) Significant toxicity is observed even in the absence of upstream BIA pathway enzymes CYP76AD5 and DODC (green lines). n=8 biological replicates per strain.

Expression of tNCS in the cytosol resulted in toxicity yet the source of this toxicity was unclear. To determine whether this toxicity results from the metabolite intermediates or from the tNCS enzyme itself, we compared the expression of tNCS in yeast with and without the upstream BIA pathway. Without the upstream enzymes CYP76AD5 and DODC, yeast cannot produce dopamine, which is one of two substrates required for (*S*)-norcoclaurine biosynthesis. Even without the upstream BIA pathway, the strain exhibited a reduced growth rate indicating that toxicity is not due to excessively high levels of (*S*)-norcoclaurine production nor is it due to the reaction between dopamine and 4-hydroxyphenylacetaldehyde (Fig. 1d). Based on this result, we concluded that the observed toxicity is not due to (*S*)-norcoclaurine biosynthesis but rather due to the tNCS protein itself, perhaps through an interaction with a native yeast factor, although the precise mechanism of this toxicity remains unknown.

### Compartmentalizing toxic tNCS inside the yeast peroxisome improves cell growth and increases titer of BIA products

Although the mechanism of tNCS toxicity is unknown, we hypothesized that we could alleviate this toxicity via subcellular compartmentalization of the tNCS protein within the cell. We reasoned that an ideal compartment would sequester the toxic tNCS enzyme away from the cytosol while allowing the free flow of NCS’s substrates and products into and out of the compartment. We identified the peroxisome as a promising candidate for subcellular compartmentalization because it can import fully-folded proteins^53^, has a high capacity for imported heterologous protein^37,44^, repurposing for compartmentalization of engineered protein should not have adverse effects on yeast viability^18,36^, and the peroxisome is permeable to small molecules^37,45^. Previous work on peroxisomes *in vitro*^45^, and our experiments on peroxisomes *in vivo*^37^, have indicated a size exclusion threshold of approximately 500-700 daltons based on the permeability of particular substrates with different molecular weights. More precisely, the permeability of small molecules across the peroxisome membrane is also affected by the molecule’s shape or ‘bulkiness,’ including its hydrodynamic radius^37,54^. We predicted that substrates dopamine and 4-hydroxyphenylacetaldehyde as well as product (*S*)-norcoclaurine would be able to cross the peroxisomal membrane given the structure and small molecular weight of these molecules (153, 136, and 271 daltons, respectively). Therefore, if tNCS were compartmentalized within the peroxisome, it would retain access to its substrates and (*S*)-norcoclaurine product would diffuse out from the peroxisome into the cytosol (Fig. 2a).

**Figure 2.**
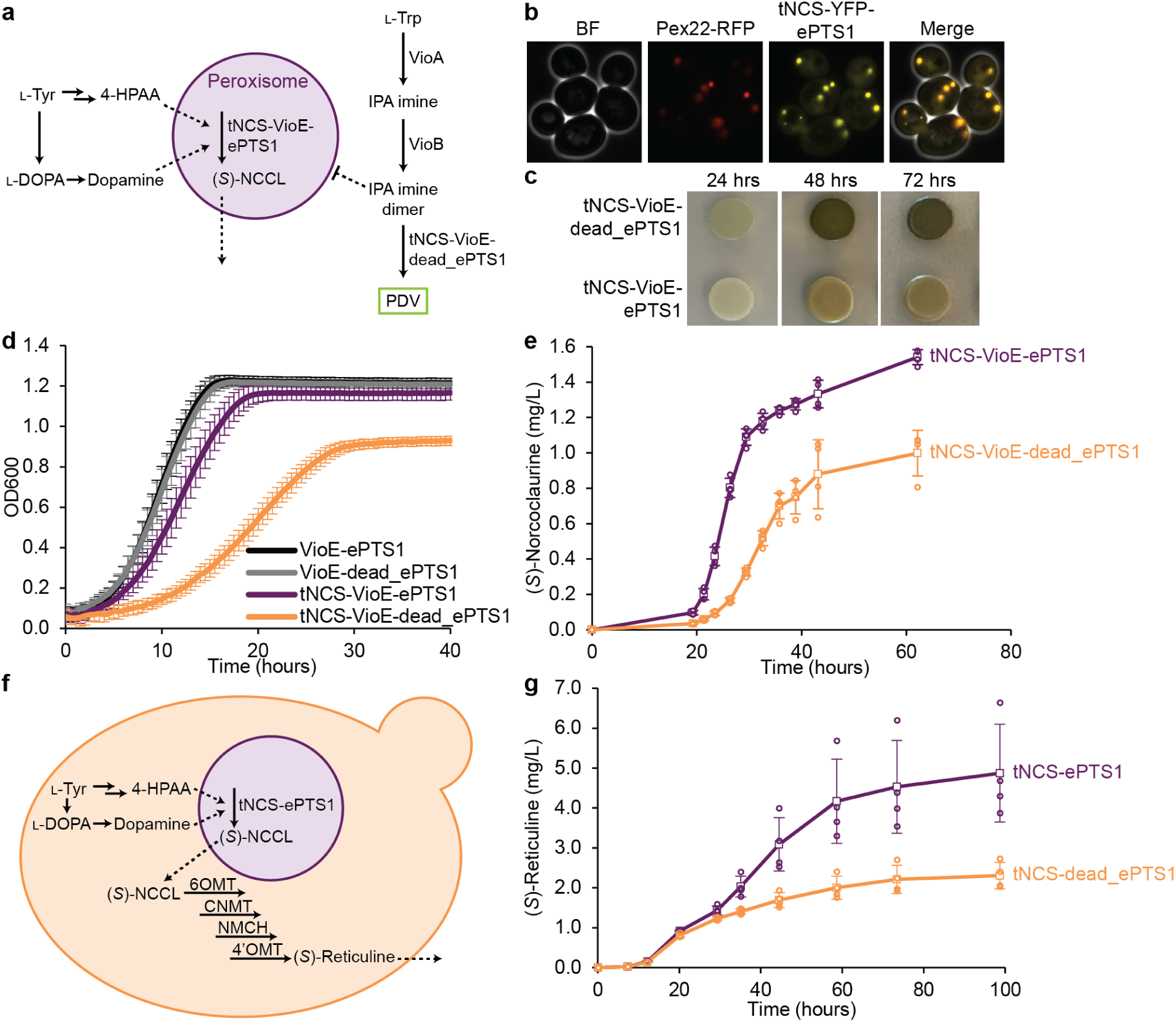
Compartmentalizing tNCS in the peroxisome alleviates cytotoxicity and increases BIA product titer. (**a**) The tNCS-VioE fusion protein will be targeted to the peroxisome when tagged with ePTS1 but will remain in the cytosol when tagged with dead_ePTS1. NCS substrates 4-hydroxyphenylacetaldehyde (4-HPAA) and dopamine can pass freely across the peroxisomal membrane to access tNCS and produce (*S*)-norcoclaurine ((*S*)-NCCL). The violacein pathway, which is used to assess peroxisomal compartmentalization, includes upstream enzymes VioA and VioB expressed in the cytosol. When VioE is also expressed in the cytosol (tNCS-VioE-dead_ePTS1), the green pigment prodeoxyviolacein (PDV) is produced. When VioE is targeted to the peroxisome (tNCS-VioE-ePTS1), less PDV is produced because the VioE substrate IPA imine dimer has low permeability across the peroxisomal membrane. (**b**) Microscopy showing co-localization of tNCS-YFP-ePTS1 with peroxisomal marker Pex22-RFP. RFP channel brightness was increased to allow visualization in the merged image. (**c**) Agar plate spots showing compartmentalization efficiency of tNCS-VioE-ePTS1 in the peroxisome compared to tNCS-VioE-dead_ePTS1. tNCS-VioE-ePTS1 spots have less green color due to lower production of PDV pigment. (**d**) Expression of tNCS in the cytosol (tNCS-VioE-dead_ePTS1, orange line) results in poor growth, whereas compartmentalization in the peroxisome (tNCS-VioE-ePTS1, purple line) alleviates much of the toxicity when compared to control strains (VioE-ePTS1 and VioE-dead_ePTS1). Error bars represent mean ± s.d. of twelve biological replicates. (**e**) Peroxisomal compartmentalization of tNCS (tNCS-VioE-ePTS1) results in higher (*S*)-norcoclaurine production compared to cytosolic expression of tNCS (tNCS-VioE-dead_ePTS1). Error bars represent mean ± s.d. of four biological replicates. (**f**) Metabolic pathway for the production of (*S*)-reticuline in *S. cerevisiae* with tNCS-ePTS1 localized in the peroxisome but with free flow of substrates, 4-HPAA and dopamine, and product, (*S*)-norcoclaurine, across the peroxisomal membrane. All other enzymes are expressed in the cytosol. (*S*)-Reticuline is natively exported from the cell and measured in the media. (**g**) Compartmentalizing tNCS-ePTS1 in the peroxisome results in higher (*S*)-reticuline production compared to tNCS-dead_ePTS1 expression in the cytosol. Error bars represent mean ± s.d. of four biological replicates.

Toxic tNCS was targeted to the peroxisome lumen using a C-terminal enhanced peroxisomal targeting signal type 1 (ePTS1). The canonical peroxisomal targeting signal consists of the amino acids SKL whereas the ePTS1 sequence includes a positively-charged amino acid linker, resulting in the C-terminal sequence LGRGRR-SKL. Compared to the canonical PTS1, use of the ePTS1 tag increases the rate of protein import into the peroxisome due to an increase in binding affinity between the tag and the cytosolic carrier protein Pex5p, which shuttles cargo to the protein import complex at the peroxisome membrane^37^. Toxic tNCS was fused to ePTS1 together with yellow fluorescent protein (YFP) to generate tNCS-YFP-ePTS1, and expressed in a yeast strain with a fluorescent peroxisomal marker consisting of the N-terminus of the peroxisomal membrane protein Pex22 fused to red fluorescent protein (RFP)^55^. Co-localization microscopy analysis of RFP and YFP signals demonstrated that ePTS1 can facilitate effective peroxisomal targeting of tNCS (Fig. 2b).

As an orthogonal assessment of peroxisomal compartmentalization, we applied a pigment-producing enzyme sequestration assay^37^ to tNCS. When three enzymes of the violacein pigment pathway, VioA, VioB, and VioE, are expressed in the cytosol, the green pigment prodeoxyviolacein (PDV) is produced. Upon fusion of ePTS1 to VioE, considerably less PDV is produced because the substrate for VioE, IPA imine dimer, has low permeability across the peroxisome membrane (Fig. 2a). Accordingly, fusions between proteins of interest and VioE-ePTS1 can provide an assessment of how well these tagged proteins are targeted to the peroxisome lumen, with better compartmentalized proteins resulting in yeast cultures with lower PDV pigment production.

The peroxisome is capable of compartmentalizing tNCS as assayed by the pigment-producing enzyme sequestration assay. We fused either our targeting tag (ePTS1) or a non-targeting tag (dead_ePTS1) to the C-terminus of tNCS-VioE to assay the efficiency of peroxisome compartmentalization. Dead_ePTS1 (LGRGRR-SKT) exchanges a threonine for the C-terminal leucine, resulting in a tag with low affinity for the cargo carrier Pex5p and, therefore, preventing import into the peroxisome^37,56^. tNCS constructs were expressed from a plasmid with a CEN6/ARS4 origin of replication and a strong pTDH3 promoter. By visually comparing the PDV pigment levels for peroxisomally targeted tNCS fused to VioE (tNCS-VioE-ePTS1) to cytosolic expression (tNCS-VioE-dead_ePTS1) at 24, 48, and 72 hours, we show that ePTS1 provides considerable compartmentalization of tNCS at these expression levels (strong pTDH3, CEN6/ARS4 copy number) (Fig. 2c). PDV was quantified by extraction from liquid culture after three days of growth and tNCS-VioE targeted to the peroxisome showed a 40% reduction in PDV pigment production relative to tNCS-VioE in the cytosol.

Production host growth and (*S*)-norcoclaurine product titer were both improved when toxic tNCS enzyme was compartmentalized in the peroxisome. In addition to the violacein pathway, our strains contained the upstream BIA pathway (CYP76AD5, DODC) so that (*S*)-norcoclaurine titer could be directly measured from the same strains. The strain with peroxisomally-targeted tNCS achieved both higher final OD and growth rate (0.103 vs 0.058 h^-1^) compared to the strain where tNCS was cytosolically expressed in small-scale growth experiments in microwell plates (Fig. 2d). In fact, the peroxisomally-targeted tNCS-VioE-ePTS1 strain grew nearly as well as our two control strains lacking toxic tNCS expression: VioE-ePTS1 (growth rate 0.136 h^-1^) and VioE-dead_ePTS1 (growth rate 0.129 h^-1^). Encouraged by these results, we conducted shake flask fermentations to compare the cell growth and (*S*)-norcoclaurine titer of tNCS-VioE-ePTS1 and tNCS-VioE-dead_ePTS1. In addition to an improvement in growth as observed in the microwell experiment (Supplementary Fig. 2), we also observed a 54% increase in final titer and a 2.2-fold increase in maximum productivity (0.13 vs 0.06 mg/L/h) (Fig. 2e).

In order to demonstrate that the (*S*)-norcoclaurine product of peroxisomally targeted tNCS is not sequestered in the peroxisome, we showed that it was accessible to the downstream four enzyme pathway that converts (*S*)-norcoclaurine to (*S*)-reticuline in the cytosol (Fig. 2f). (*S*)-Reticuline represents the last shared intermediate of the large and highly branched family of BIA natural products^29^. In order to synthesize (*S*)-reticuline, (*S*)-norcoclaurine must undergo two *O*-methylations, one *N*-methylation, and one hydroxylation, catalyzed by 6-*O*-methyltransferase (6OMT), coclaurine *N*-methyltransferase (CNMT), *N*-methylcoclaurine 3’-hydroxylase (NMCH), and 4’-*O*-methyltransferase (4’OMT)^29^. All four of these downstream enzymes were cytosolically expressed along with the upstream tyrosine hydroxylase CYP76AD5 and L-DOPA carboxylase DODC with NCS compartmentalized in either the peroxisome or cytosol. Compared to cytosolic expression (dead_ePTS1), peroxisomal targeting (ePTS1) resulted in both a modest growth rate improvement and a 34% increase in final OD (Supplementary Fig. 3). A corresponding titer improvement was also observed (Fig. 2g), with a 2.1-fold increase in final titer and a 34% increase in maximum productivity (0.11 vs 0.08 mg/L/h). In general, (*S*)-reticuline titers were higher than (*S*)-norcoclaurine titers. Cell density was also higher for (*S*)-reticuline strains compared to (*S*)-norcoclaurine strains (Supplementary Fig. 2 vs Supplementary Fig. 3). We attribute these differences to the inclusion of enzymes from the violacein pathway (VioA, VioB, and tNCS fused to VioE) in the (*S*)-norcoclaurine production strains, whereas the (*S*)-reticuline producing strains do not contain any of the violacein pathway enzymes.

### Incomplete compartmentalization occurs at very high expression levels of tNCS

While compartmentalization of tNCS in the peroxisome is effective at the tested expression levels, we wanted to determine if the peroxisome is capable of compartmentalizing even higher expression levels of tNCS to achieve increased titers. To characterize the upper limits of enzyme targeting, we investigated peroxisome targeting when the plasmid copy number was increased from CEN6/ARS4 (2-5 copies per cell) to 2-micron (2µ) (15-50 copies per cell)^50,57^. At this very high expression level, peroxisomal compartmentalization was still effective but incomplete (Fig. 3). At high 2µ expression level, much of the tNCS-YFP-ePTS1 protein was targeted to the peroxisome as visualized by microscopy using Pex22-RFP as a peroxisomal marker (Fig. 3a). However, more residual cytosolic tNCS-YFP-ePTS1 localization was observed at this higher expression level, indicated by increased cytosolic YFP signal compared to the previous CEN6/ARS4 expression (Fig. 3a vs Fig. 2b). At 2µ expression levels, incomplete compartmentalization and high toxicity were observed in the pigment-producing enzyme sequestration assay and in growth experiments. PDV pigment extraction after three days of growth showed a 39% reduction in PDV pigment production when tNCS-VioE was targeted to the peroxisome instead of the cytosol (Fig. 3b). Toxicity due to high cytosolic expression of tNCS-VioE was apparent in the agar plate spots, evidenced by low cell density at 24 hours and abnormal spot morphology (wrinkled appearance) at 48 and 72 hours. Growth curves showed that cytosolic expression (tNCS-VioE-dead_ePTS1) was highly toxic, with hardly any measurable growth occurring before 20 hours, at which point the control strains lacking tNCS, VioE-ePTS1 and VioE-dead_ePTS1, had already reached a saturated cell density (Fig. 3c). Peroxisomal compartmentalization of tNCS (tNCS-VioE-ePTS1) was able to alleviate some of the toxicity, confirming that peroxisome targeting was still effective; however, final OD and growth rate were below that of the control strains, suggesting the peroxisome is not natively capable of completely compartmentalizing the toxic tNCS at these elevated expression levels.

**Figure 3.**
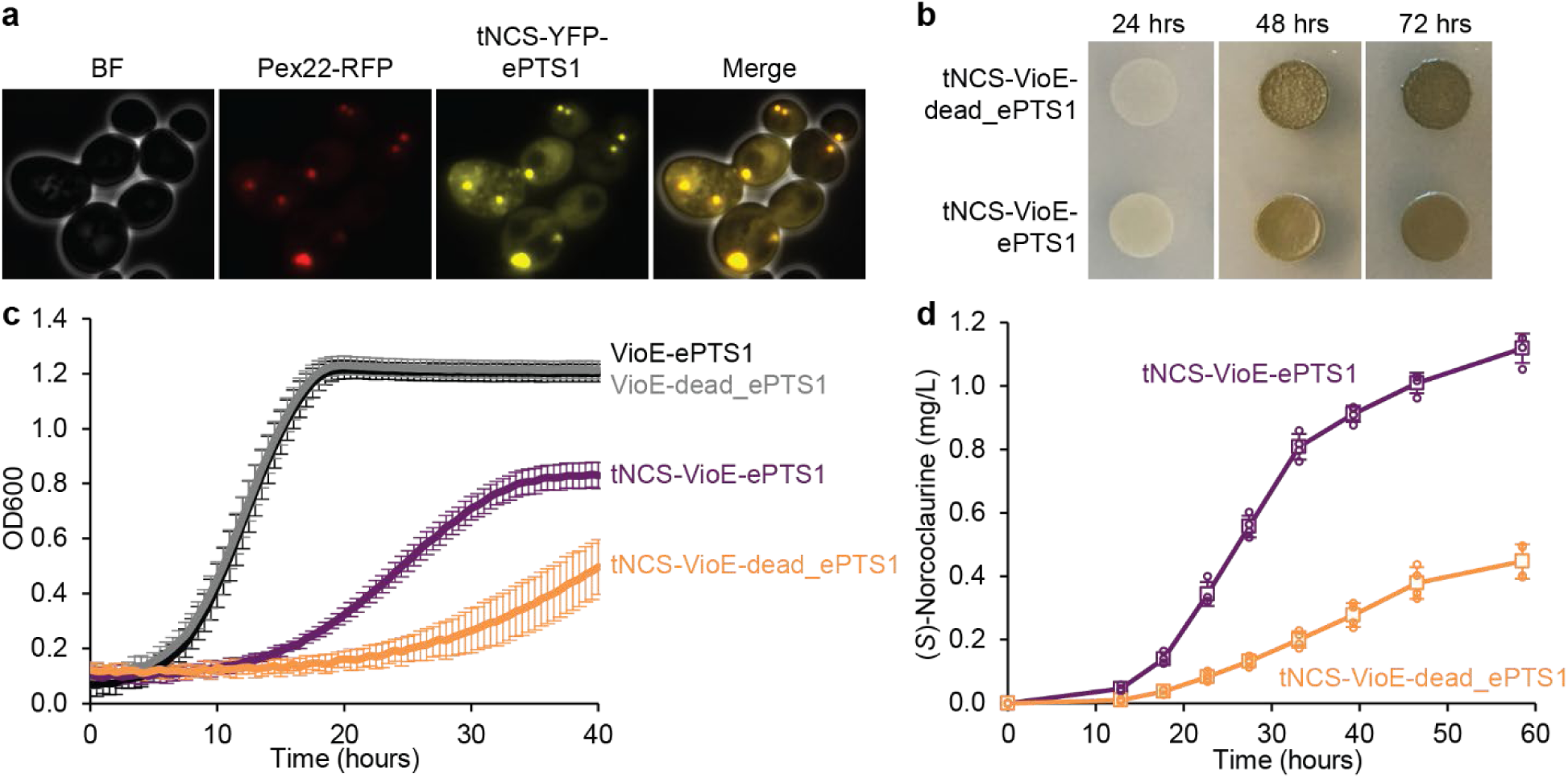
Peroxisomal targeting of toxic tNCS at a very high expression level improves growth and (*S*)-norcoclaurine titer, but compartmentalization is incomplete. (**a**) Microscopy showing co-localization of tNCS-YFP-ePTS1 (pTDH3 2μ expression level) with Pex22-RFP at peroxisomes and residual YFP signal in the cytosol (incomplete compartmentalization). RFP channel brightness was increased to allow visualization in the merged image. (**b**) Agar plate spots showing compartmentalization efficiency of tNCS-VioE-ePTS1 in the peroxisome compared to tNCS-VioE-dead_ePTS1. Abnormal spot morphology for cytosolic expression (tNCS-VioE-dead_ePTS1) indicates high toxicity. (**c**) Expression of tNCS in the cytosol (tNCS-VioE-dead_ePTS1) results in very poor growth (orange line) whereas peroxisomal expression (tNCS-VioE-ePTS1) partially alleviates toxicity at pTDH3 2μ expression level (purple line) when compared to growth of control strains (black and grey lines). Error bars represent mean ± s.d. of twelve biological replicates. (**d**) Peroxisomal targeting of tNCS (tNCS-VioE-ePTS1) results in higher (*S*)-norcoclaurine production compared to cytosolic expression of tNCS (tNCS-VioE-dead_ePTS1). Error bars represent mean ± s.d. of four biological replicates.

Production of (*S*)-norcoclaurine was greatly improved upon targeting of tNCS to the peroxisome compared to the cytosol at 2µ expression level. Final (*S*)-norcoclaurine titer is an impressive 2.5-fold higher for peroxisomal expression compared to cytosolic expression from a 2µ plasmid (Fig. 3d), and maximum productivity is 3.2-fold higher (0.045 vs 0.014 mg/L/h). However, we observed that the final titer of peroxisomal expression from a 2µ plasmid is lower than that of peroxisomal CEN6/ARS4 expression (1.1 vs 1.5 mg/L). We attribute this lower titer to the poor growth caused by incomplete compartmentalization at the elevated expression level of the 2µ plasmid. Indeed, the final OD and growth rate in flasks are 3.6 and 0.34 h^-1^, respectively, when tNCS is expressed from a CEN6/ARS4 plasmid while only 2.7 and 0.14 h^-1^ when expressed from a 2µ plasmid (Supplementary Fig. 2 vs Supplementary Fig. 4).

### Developing strategies to improve compartmentalization capacity of peroxisome-targeted cargo

Protein capacity is limited in most organelles and, although the peroxisome can natively compartmentalize a comparably impressive amount^44^, further increased capacity is desired for many applications, including the insulation of the toxic tNCS enzyme. In *Saccharomyces cerevisiae*, the natural function of peroxisomes is the β-oxidation of long-chain fatty acids^58^. In glucose-containing media, peroxisomes have an average diameter of 0.2 µm^59^, but in media containing the C18 fatty acid oleate, peroxisomes are enlarged to diameters of 0.3-0.5 µm^60,61^. Glucose and other sugars repress the peroxisome induction effect, but growth on oleate as the sole carbon source is poor compared to growth on glucose^62^. Accordingly, induction media commonly uses an alternative carbon source such as glycerol in addition to oleate^63^. We found that yeast growth was also poor in this induction media (as has been reported for *S. cerevisiae* in glycerol^64^). To balance growth rate with the ability to induce higher peroxisome capacity, we developed a hybrid induction media that contained 0.5% glucose in addition to 10% glycerol and 0.1% oleate (compared to standard media containing 2% glucose). Given that glucose will be preferentially consumed over other carbon sources in *S. cerevisiae*, we reasoned that 0.5% glucose would support fast initial culture growth before being fully utilized. This initial biomass would subsequently switch to consumption of glycerol, thereby permitting proliferation of large peroxisomes due to oleate induction.

Peroxisomes appeared brighter in YFP fluorescence when induced in oleate growth media and visualized by peroxisome-targeted YFP and microscopy (Supplementary Fig. 5). In order to capture and compare the relative differences in YFP fluorescence when cells were grown in different media, a fairly low exposure setting was used for all microscopy images to avoid overexposure of YFP signal in oleate media. As a result, YFP fluorescence of peroxisomes grown in only glucose look very dim in comparison, although YFP targeting was confirmed both using higher exposure settings during image capture and by post image processing. Both the standard oleate glycerol induction media and our hybrid induction media (glucose, glycerol, and oleate) showed brighter YFP peroxisome puncta compared to cells grown in glucose only, which we interpret as being consistent with oleate-induced peroxisome proliferation to accommodate increased targeting of fatty acid utilization enzymes. We then performed a (*S*)-norcoclaurine production experiment using toxic tNCS targeted to the peroxisome and expressed on the high copy 2µ plasmid, comparing standard glucose media to our hybrid induction media (Supplementary Fig. 6). Growth and final cell density were decreased in the hybrid induction media compared to standard glucose media. Consequently, final (*S*)-norcoclaurine titer was reduced 14% in the hybrid induction media; however, normalizing titer by OD showed a 21% *higher* production, indicating higher per cell productivity and suggesting that peroxisome induction could be a promising strategy for increasing protein cargo capacity if growth were further improved.

To avoid the poor growth phenotype that results from use of glycerol as the main carbon source with oleate induction, we sought to engineer a strain with increased peroxisome capacity while grown on the favored glucose carbon source by constitutively expressing transcription factors involved in peroxisome proliferation. We focused on three transcription factors known to control peroxisome proliferation - ADR1, OAF1, and PIP2^65–67^. All three transcription factors are post-translationally repressed by glucose^68,69^ and OAF1 is post-translationally induced by oleate^70^. We aimed to make ADR1, OAF1, and PIP2 constitutively active in standard growth media containing glucose as well as remove the requirement for oleate by engineering or deleting the regulatory domains of these transcription factors.

An engineered version of the ADR1 transcription factor was designed to enable constitutive activation in glucose media. ADR1 is known to be inactivated by phosphorylation at Ser-230 under high glucose concentrations^71^. ADR1 activation is achieved by Ser-230 dephosphorylation when glucose concentration is low. Mutation of Ser-230 to alanine has been shown to mimic the dephosphorylated, active state, leading to transcription of downstream genes even in the presence of glucose^72^. We applied this S230A mutation to generate ADR1c, a constitutively active version of ADR1.

Engineered versions of OAF1 and PIP2 transcription factors were designed through removal of regulatory domains to enable activation in glucose media without oleate. OAF1 and PIP2 are paralogs with similar domain architectures, both containing an N-terminal DNA binding domain, followed by regulatory domains, and ending with a C-terminal activation domain^69^. We hypothesized that removal of the regulatory domains would enable constitutive activation in standard growth media by removing glucose repression and the need for oleate induction. For OAF1, we specifically tethered amino acids 1-100 (containing the DNA binding domain) via a flexible two amino acid glycine-serine linker to amino acids 807-1047 (containing the activation domain). Likewise, for PIP2 we tethered amino acids 1-56 (containing the DNA binding domain) to amino acids 833-996 (containing the activation domain) via a two amino acid glycine-serine linker. We called these engineered transcription factors OAF1c and PIP2c, respectively.

A fluorescence-based assay was devised to test peroxisome compartmentalization capacity upon expression of combinations of the three engineered transcription factors (ADR1c, OAF1c, PIP2c). The assay utilizes YFP fused to an N-terminal proteasome-based degradation signal, UbiY^52,73^, and a C-terminal ePTS1 tag (Fig. 4a). In this assay, Pex5p recognizes the ePTS1 tag and shuttles UbiY-YFP-ePTS1 cargo to the peroxisomal membrane for import where it would be protected from degradation. Residual UbiY-YFP-ePTS1 remaining in the cytosol is recognized and targeted to the proteasome for degradation. The fluorescent signal should, consequently, only reflect YFP localized in the peroxisome. Therefore, combinations of constitutively-active transcription factors resulting in improved cargo capacities will be reflected as increased YFP signal compared to the signal produced in a strain with no transcription factor overexpression.

**Figure 4.**
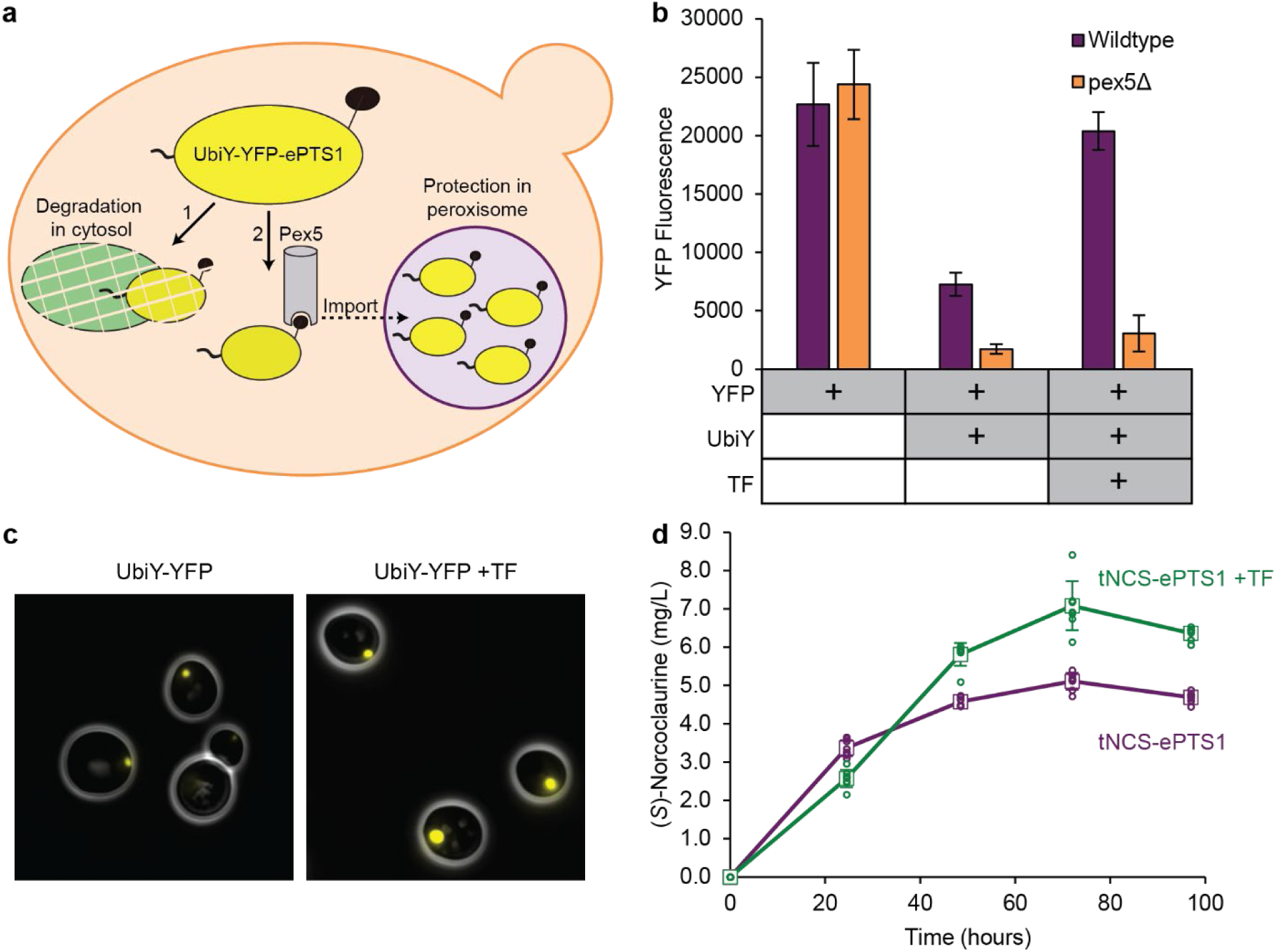
Genetic induction of peroxisome proliferation can address incomplete compartmentalization at very high expression levels of tNCS. (**a**) Degron-YFP assay for measuring peroxisomal compartmentalization. Residual UbiY-YFP-ePTS1 that is not imported to the peroxisome by Pex5p (path 2) is degraded in the cytosol by the proteasome (path 1). Changes in YFP signal are therefore indicative of changes in peroxisome import capacity. (**b**) Degron-YFP assay was used to assess transcription factor overexpression. Strains with UbiY-YFP-ePTS1 expressed in the cytosol (pex5Δ) have lower fluorescence than their corresponding peroxisomally-targeted strains (Wildtype) due to UbiY-YFP degradation by the proteasome. TF represents expression of engineered transcription factors ADR1c, OAF1c, and PIP2c. Error bars represent mean ± s.d. of twelve biological replicates. (**c**) Microscopy of UbiY-YFP-ePTS1 cells shows larger peroxisomes when ADR1c, OAF1c, and PIP2c are expressed (+TF). YFP channel brightness was increased identically across both images to allow better visualization of peroxisomes from the UbiY-YFP strain. Full images showing population variability are available in Supplementary Fig. 9. (**d**) (*S*)-Norcoclaurine production is improved upon expression of engineered transcription factors. Error bars represent mean ± s.d. of eight biological replicates.

The degron-YFP assay functioned as expected, with degradation observed upon cytosolic expression and protection observed upon peroxisomal targeting. We demonstrated that fusion of the UbiY degron to YFP-ePTS1 resulted in cytosolic degradation by expressing this fusion in a pex5 knockout strain (pex5Δ), which prevents peroxisomal import (Fig. 4b, second orange bar). Expression of this fusion protein, UbiY-YFP-ePTS1, in a wildtype background strain containing Pex5p resulted in 4.2-fold higher YFP signal (Fig. 4b, second purple bar), indicating protection from degradation due to peroxisomal import. Although considerable protection was achieved, this signal was 32% of the YFP signal measured for a strain containing YFP without the UbiY degron (Fig. 4b, first purple bar), indicating that compartmentalization is not complete.

Expression of engineered transcription factors increased protection of cargo from degradation (Fig. 4b) and resulted in visibly larger peroxisomes with increased fluorescence from importing UbiY degron-fused YFP (Fig. 4c). The engineered transcription factors ADR1c, OAF1c, and PIP2c were first expressed in different combinations and assayed for UbiY-YFP-ePTS1 signal after growth for 48 hours in a 96-well culture block (Supplementary Fig. 7). The highest YFP signal, indicating improved protection from degradation, was observed upon expression of all three engineered transcription factors. We then investigated the dynamics of the system by directly culturing the triple-transcription factor strain (+TF), alongside controls, in a small-scale microwell plate (Supplementary Fig. 8). Strains with UbiY-YFP-ePTS1 expressed in the cytosol showed an increase in YFP signal over the first ∼15 hours during the culture growth phase, and thereafter showed a drop in YFP signal as the rate of proteasomal degradation outpaced the rate of YFP synthesis, as expected. Strains with UbiY-YFP-ePTS1 targeted to the peroxisome exhibited protection from degradation by showing less YFP degradation after the growth phase, resulting in higher YFP signal at the endpoint relative to their cytosolic counterparts. Endpoint values from the timecourse experiment are depicted in Fig. 4b and show that the triple-transcription factor combination (TF) resulted in 2.8-fold higher UbiY-YFP-ePTS1 fluorescence than without TF expression and 90% of the fluorescence compared to the no-degron strain. We then compared the wildtype background strain to the ADR1c/OAF1c/PIP2c expression strain, both expressing UbiY-YFP-ePTS1, by microscopy and found that transcription factor overexpression resulted in larger peroxisomes (Fig. 4c). While there was cell-to-cell variability with regard to the presence and size of peroxisomes, they were routinely larger and brighter in the +TF strain (Supplementary Fig. 9). Thus, increased cargo capacity can be achieved in glucose media through the expression of constitutively-active versions of transcription factors involved in glucose derepression and oleate induction.

Improved capacity for UbiY-YFP-ePTS1 translated to a 36% improvement in (*S*)-norcoclaurine titer when tNCS is targeted to the peroxisome compared to peroxisome targeting without transcription factor overexpression. We built strains containing the high copy 2µ plasmid expressing peroxisome-targeted tNCS (pTDH3-tNCS-ePTS1), in either a wildtype strain background or a strain containing the best set of transcription factors overexpressed (ADR1c/OAF1c/PIP2c). Transcription factor overexpression resulted in a 36% increase in (*S*)-norcoclaurine titer compared to the wildtype strain without transcription factor overexpression (Fig. 4d). Although we did not observe the expected increase in growth upon expression of constitutively active ADR1c/OAF1c/PIP2c (Supplementary Fig. 10), we also noted that transcription factor overexpression resulted in a decrease in growth even in the absence of toxic tNCS (Supplementary Fig. 11). Thus, one or more of the over 100 genes regulated by these transcription factors presents a burden or directly inhibits growth^74,75^. When normalized by OD, expression of the engineered transcription factors resulted in a 47% improvement in (*S*)-norcoclaurine production (Supplementary Fig. 10). Ultimately, the benefit of transcription factor overexpression on peroxisome size and compartmentalization capability outweighed the growth defect, resulting in a higher final titer of (*S*)-norcoclaurine.

## Discussion

Toxicity of heterologous proteins^76–82^ and metabolites^83–89^ is a significant problem in metabolic engineering. Toxicity manifests as substantial obstructions to large scale fermentation in terms of slower growth of cells and ultimately lower titer and productivity. Our group faced the challenge of needing to highly express tNCS enzyme in order to improve titer of BIAs while discovering that high expression of tNCS also caused cytotoxicity in *S. cerevisiae*. With regard to compartmentalization, this challenge required a specific solution, one in which the tNCS protein would be efficiently targeted and sequestered but retain access to its small molecule substrates and allow the (*S*)-norcoclaurine product to exit to the cytosol where the upstream and downstream enzymes are localized. One approach could involve the use of a compartment that is not readily permeable to small molecules (e.g., vacuole^90^) and compartmentalization of the entire metabolic pathway. However, in addition to the requirement for high protein capacity that has not been demonstrated for most organelles, this approach would necessitate the incorporation of one or several transporters recognizing the enzyme’s substrate(s) and, potentially, product at the organelle membrane.

Peroxisomal compartmentalization has the potential to be a generalizable strategy for expression of toxic proteins. As is the case for many engineered metabolic pathways that suffer from toxicity, the mechanism of tNCS toxicity is not known. The mechanisms causing toxicity can be difficult to elucidate. A common strategy for minimizing, or even overcoming, this undesired effect on the production host is to conduct adaptive laboratory evolution (ALE)^91^. ALE works best when the desired flux can be selected such that derived mutations for improved growth are not exclusive with desired pathway flux, and ALE can fail if growth is improved at the expense of product titer^92^. ALE often requires months of cell passaging to enable mutations to be acquired, especially when the acquisition of multiple mutations are required^91^. We conducted ALE on a strain expressing tNCS followed by a secondary screen of the fastest growing strains for (*S*)-norcoclaurine titer. The first type that we observed, a strain that downregulates or loses entirely the expression of tNCS, is detrimental to product formation. The second type that we observed was a strain that appeared to alleviate toxicity while producing high (*S*)-norcoclaurine titer, comparable to the titer from our 2µ plasmid-expressed tNCS-ePTS1 strain. Unfortunately, sequencing the genome of this adapted strain did not yield any decipherable mutations to explain how the toxicity was alleviated. Although frequently successful, ALE will not work for all toxic proteins. If the mechanism of toxicity is well understood, a more rational solution may be developed, such as a mutation to the protein to block undesired interactions, or a genomic modification (gene deletion or mutation) that reduces toxicity. Compartmentalization in the peroxisome is a considerably more rapid strategy to employ, requiring only the addition of the C-terminal ePTS1 tag and does not require understanding of the mechanism of toxicity other than it is due to the protein itself and not any metabolite product. Further, this strategy should generalize to all soluble proteins directly exerting toxicity (i.e. via an interaction with protein(s), lipid(s), carbohydrate(s), etc. accessible in the cytosol).

The peroxisome in *S. cerevisiae* has evolved for the compartmentalization of increased copies of enzymes required for the β-oxidation of long-chain fatty acids^58^. Examples in nature indicate that the peroxisome is an inherently flexible organelle and, therefore, likely has the potential to be engineered as a synthetic compartment for desired applications. In addition to functioning as a catabolic compartment for the β-oxidation pathway, peroxisomes are used for unique biochemical reactions, such as penicillin biosynthesis^93^ and generation of light in fireflies^94^. Moreover, peroxisome size, number, and morphology can be substantially altered in different organisms using a variety of substrates such as oleate^60^, methanol^95,96^, D-alanine^97^, and butyrate^98^ indicating that genetic circuits for peroxisome proliferation are prevalent across a broad range of eukaryotes. We pursued genetic reprogramming of peroxisome proliferation via constitutive expression of engineered transcription factors and demonstrated increased cargo capacity without the requirement for oleate. While this is an important step toward genetic control over peroxisome cargo capacity, expression of these constitutively-active transcription factors resulted in a decrease in growth even in the absence of toxic tNCS. This should not be too surprising given that approximately 100 downstream genes are regulated by these transcription factors and, accordingly, one or more are likely to have an impact on cell state/growth^74,75^. Future work will seek to identify the critical downstream targets of ADR1, OAF1, and PIP2 such that only those critical genes will be manipulated to control cargo capacity without impeding growth as was observed when the engineered transcription factors were overexpressed.

Thus, this work establishes a robust strategy for circumventing the growth defect resulting from the expression of a cytotoxic metabolic enzyme while enabling increased flux. Protection of heterologous cargo from the effects of the host cell, such as proteasome-based degradation, was also demonstrated. Accordingly, peroxisome compartmentalization offers the engineer bidirectional benefits: protection of the host cell from the heterologous protein as well as protection of the heterologous protein from the host cell. This work highlights the peroxisome as a versatile, engineerable organelle that can be utilized to address challenges in metabolic engineering.

## Methods

### Strains and growth media

The base *S. cerevisiae* strain for all experiments was BY4741 (*MATa his3Δ1 leu2Δ0 met15Δ0 ura3Δ0*). Base strain BY4741 and BY4741 *pex5Δ* were ordered from Open Biosystems—GE Dharmacon. Wildtype yeast cultures were grown in YPD (10 g/L Bacto yeast extract; 20 g/L Bacto peptone; 20 g/L glucose). Selection of auxotrophic markers (URA3, LEU2, HIS3) was performed in synthetic complete (SC) media (6.7 g/L Difco Yeast Nitrogen Base without amino acids (Spectrum Chemical); 2 g/L Drop-out Mix Synthetic minus appropriate amino acids, without Yeast Nitrogen Base (US Biological); 20 g/L glucose). All strains used in this work are listed in Supplementary Table 1.

Oleate induction media was prepared as 6.7 g/L Difco Yeast Nitrogen Base without amino acids (Spectrum Chemical); 2 g/L Drop-out Mix Synthetic minus appropriate amino acids, without Yeast Nitrogen Base (US Biological), with the following variations (final concentrations described): 1) 2% glucose, 2) 10% glycerol, 0.1% oleic acid, 0.4% Tween80, 3) 0.5% glucose, 10% glycerol, 0.1% oleic acid, 0.4% Tween80.

Golden gate assembly reactions were transformed into chemically competent *Escherichia coli* prepared from strain TG1 (Lucigen). Transformed cells were selected on Lysogeny Broth containing the antibiotics chloramphenicol, ampicillin, or kanamycin.

### Yeast strain construction

Yeast expression vectors were built using Golden Gate Assembly as described in the YTK system^52^. Integration into the yeast genome via homologous recombination at the URA3 or LEU2 locus was achieved by transformation of linearized plasmids (NotI digestion, NEB) whereas replicating CEN6/ARS4 or 2µ plasmids were transformed directly into yeast without pre-digestion with NotI. All transformations were performed using a standard lithium acetate method^99^ and cells were plated onto selective auxotrophic SC 2% glucose agar plates. All strains are described in Supplementary Table 1. Individual colonies (biological replicates) were picked directly from this transformation plate and grown independently for further analysis.

Chromosomal integrations of Pex22-RFP, VioA, and VioB were performed by co-transforming a CEN6/ARS4 CRISPR plasmid (containing Cas9, guide RNA for targeting the appropriate locus, HIS3 auxotrophic marker) with linearized repair DNA designed to integrate the gene(s) of interest. Cells were plated on histidine dropout media, restreaked on histidine dropout media, then grown in non-selective media to remove the CRISPR plasmid. Chromosomal integration was confirmed by PCR and removal of CRISPR plasmid was confirmed by replica plating from non-selective media onto histidine dropout media (colony will not grow on histidine dropout media if CRISPR plasmid has been removed).

All plasmids are available upon request from the authors.

### Growth curves in microwell plates

Individual colonies were grown to saturation in SC 2% glucose media with auxotrophic selection at 30°C with shaking. Saturated cultures were then diluted 50-fold in fresh selective media and grown for 6-7 hours before a second dilution (10-fold) in 96-well Costar microplates (black, clear bottom) and sealed with breathe-easy film (Sigma). OD600 was measured at 30 minute intervals using a microplate reader (Tecan Spark), with continuous orbital shaking at 30°C in a humidity cassette. Variation between biological replicates was calculated as standard deviation of the mean OD600 at each time point and represented as error bars in figures.

### Fluorescence microscopy

For confirmation of protein localization, strains were grown to saturation in SC 2% glucose media with auxotrophic selection, diluted 50-fold into fresh selective media and grown for a further 6-8 hours. Cultures were resuspended in 1x PBS before spotting onto plain glass slides for imaging on a Zeiss Axio observer D1 microscope with X-Cite Series 120 fluorescent lamp and Hamamatsu Orca-Flash 4.0 Digital Camera. Images were analyzed using ZEN 3.0 (blue edition) software (Zeiss). Fluorescent protein variants used in this study were the yellow fluorescent protein Venus and the red fluorescent protein mRuby2.

### Spot assays for PDV production and PDV quantification

Agar plate spots for visualization of PDV production were generated by plating 5 µl of saturated culture on SC 2% glucose agar plates with appropriate auxotrophic selection. Plates were grown at 30°C and imaged at 24, 48 and 72 hours. For PDV quantification, strains were grown to saturation at 30°C with shaking in SC 2% glucose media with auxotrophic selection. Saturated cultures were then diluted 50-fold into fresh selective media and grown for a further 72 hours before PDV extraction was performed as previously described^37^. Relative PDV quantification was determined using bulk fluorescence measurements with 100 μl extracts on a microplate reader (Tecan Infinite M1000 Pro) at excitation 535/5 nm and emission 585/5 nm. PDV production can be estimated using fluorescence measurements at these wavelengths due to a linear correlation to extracts quantified by HPLC^37^. A minimum of six biological replicates of each strain was used for PDV quantification.

### BIA production in 96-well culture blocks

(*S*)-Norcoclaurine production experiments for Fig. 1b, Fig. 4d, Supplementary Fig. 1, Supplementary Fig. 6, and Supplementary Fig. 10 were performed in 96-well culture blocks. Individual colonies were grown to saturation at 30°C in selective media before 50-fold dilution into fresh selective media. For Fig. 1b and Supplementary Fig. 1, cultures were grown for 72 hours after the 50-fold dilution. For Supplementary Fig. 6, cultures were grown for 160 hours after the 50-fold dilution. For Fig. 4d and Supplementary Fig. 10, the inoculation culture was diluted 50-fold into a separate block for each timepoint. At each timepoint or at final harvest, samples were taken for measurement of OD600 on a microplate reader (Tecan Infinite M1000 Pro). The remaining culture was centrifuged at ∼4000 g and the supernatant filtered with an Acroprep Advance 0.2 µl filter plate (Pall Corporation). Eight-point standard curves using (*S*)-norcoclaurine chemical standard (Bepharm) were generated as two-fold serial dilutions prepared in culture media and used to convert LC/MS peak area to concentration. Samples and standards were diluted in water prior to LC/MS analysis to achieve concentrations within the linear range of detection of the instrument. Data shown is the mean (*S*)-norcoclaurine titer for three-to-eight biological replicates of each strain (as indicated in each figure legend) and error bars represent standard deviation of the mean.

### BIA production in shake flasks

Shake flask fermentation experiments were performed for (*S*)-norcoclaurine production with tNCS expressed at CEN6/ARS4 level (Fig. 2e, Supplementary Fig. 2), (*S*)-reticuline production with tNCS at CEN6/ARS4 level (Fig. 2g, Supplementary Fig. 3), and (*S*)-norcoclaurine with tNCS at 2µ level (Fig. 3d, Supplementary Fig. 4). Individual colonies were grown to saturation at 30°C in selective media. For Fig. 2e and Supplementary Fig. 2, saturated cultures were diluted 500-fold into 50mL selective media in 250mL baffled shake flasks. For all other experiments, saturated cultures were diluted to a starting OD of 0.05 in 50mL selective media in 250mL baffled shake flasks. Shake flask cultures were grown at 30°C with shaking at 220 rpm in an Innova 44 incubator (Eppendorf). At each timepoint, samples were taken for measurement of OD600 (Thermo Scientific Genesys 30 spectrophotometer) and BIA production. BIA samples were processed as described above for LC/MS and compared to an (*S*)-norcoclaurine or (*S*)-reticuline (ChemCruz) standard curve for quantification.

### Fluorescence measurements for degron-YFP assay

Individual colonies were grown to saturation in SC 2% glucose media with auxotrophic selection at 30°C with shaking. Saturated cultures were then diluted 50-fold in fresh media to a final volume of 500 µL in Corning 96-well culture blocks (Catalog number 07-200-700) and incubated at 30°C with shaking at 750 rpm. To sample OD600 and YFP fluorescence, 100 µL was aliquoted into 96-well Costar microplates (black, clear bottom) and measured on a Tecan Infinite M1000 Pro plate reader. YFP fluorescence was measured using an excitation wavelength of 515 nm and emission wavelength of 528 nm (5 nm bandwidth for excitation and emission). Variation between biological replicates was calculated as standard deviation of the mean and represented as error bars in figures.

Measurement of the dynamics of degron-YFP was performed as above for growth curves in microwell plates, but included the measurement of YFP in addition to OD600 at every timepoint. YFP fluorescence was measured using an excitation wavelength of 513 nm (5 nm bandwidth) and an emission wavelength of 531 nm (7.5 nm bandwidth).

### LC/MS analysis

Liquid chromatography/mass spectrometry (LC/MS) was performed using a 1260 Infinity LC System connected to a 6120 Quadrupole Mass Spectrometer (Agilent Technologies). All samples and standards were diluted in water prior to injection, to achieve linear calibration curves for (*S*)-norcoclaurine and (*S*)-reticuline with all samples falling within the bounds of the calibration curve. Ten microliters of each diluted sample were injected and sample separation was achieved using a Zorbax Eclipse Plus C18 guard column (4.6 cm×12.5 cm, 5 μm packing, Agilent Technologies) connected to a Zorbax Eclipse Plus C18 column (4.6 mm × 100 mm, 3.5 μm packing, Agilent Technologies) at ambient temperature using a 0.5 mL/min flow rate. Samples were eluted with a linear gradient from 100% water/0% acetonitrile plus 0.1% formic acid to 65% water/35% acetonitrile plus 0.1% formic acid over the course of 15 min. MS was conducted in atmospheric pressure ionization-positive electrospray (API-ES positive) mode at 100-V fragmentor voltage with ion detection set to targeted detection of (*S*)-norcoclaurine (272.1 m/z) and/or (*S*)-reticuline (330.2 m/z).

## Data Availability

The datasets generated and analyzed during the current study are available from the corresponding author on reasonable request. Accession codes for plasmid sequences will be available before publication.

## Acknowledgements

We thank members of the Dueber Lab for valuable assistance and feedback throughout this project, particularly Zachary Russ for training and Kay Siu for useful discussions; Sunnyjoy Dupuis for construction of the Pex22-RFP strain; and members of the Wenjun Zhang lab for assistance with LC/MS, particularly Will Skyrud and Antonio Del Rio Flores. This work is supported by NSF grant MCB 1818307 and by the Center for Cellular Construction, an NSF Science and Technology Center, under grant agreement DBI-1548297.

## Author Contributions

P.S.G., J.A.S., J.J.B., and J.E.D. designed the research. P.S.G., J.A.S., J.J.B., and B.C. performed the experiments. P.S.G., J.A.S., and J.J.B. analyzed the results. J.E.D. supervised the research. P.S.G., J.A.S., J.J.B., and J.E.D. wrote the manuscript.

