## Supplementary Info for "Repurposing the yeast peroxisome to compartmentalize a toxic enzyme enables improved (*S*)-reticuline production"

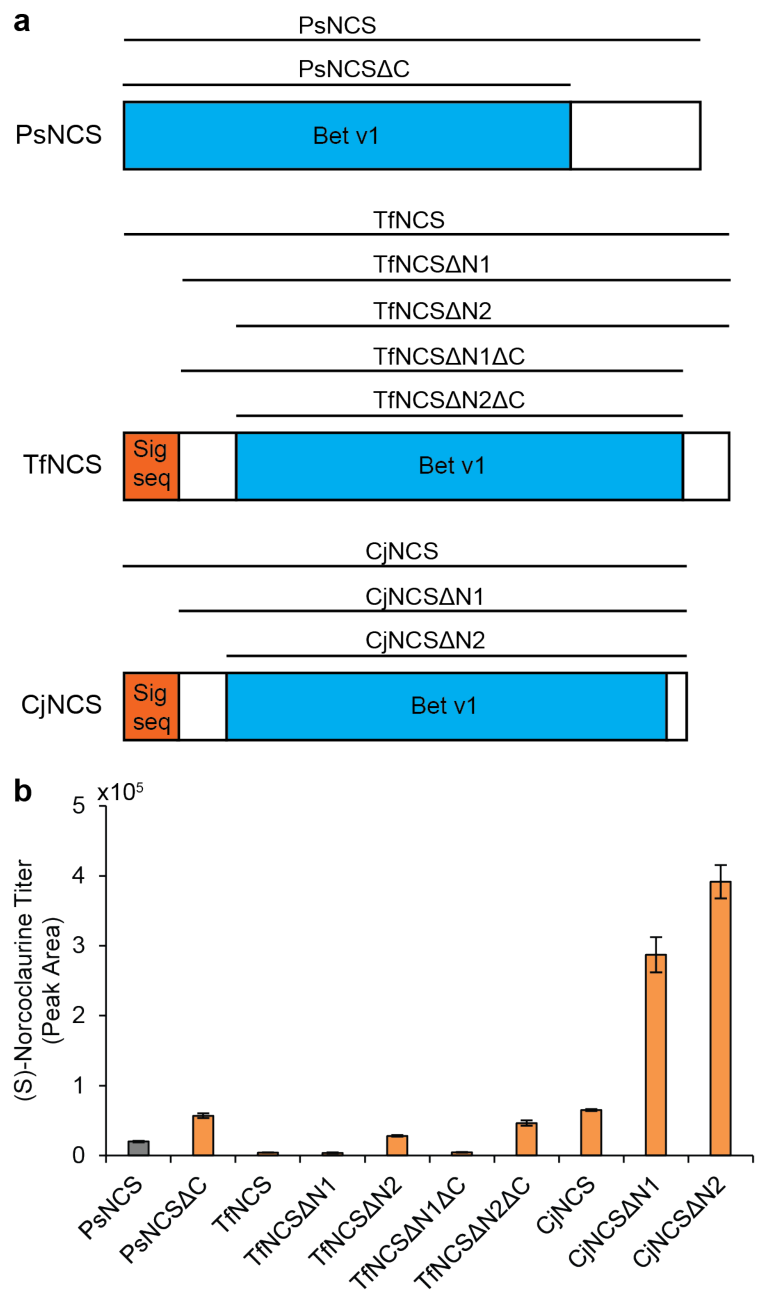


**Supplementary Figure 1. NCS homologues and truncated variants.** (**a**) Truncation strategy for NCS homologues from *Papaver somniferum* (Ps), *Thalictrum flavum* (Tf), and *Coptis japonica* (Cj). Signal sequence (Sig seq) and Bet v1 domains were identified using the SMART domain analysis tool[1]. (**b**) (*S*)-Norcoclaurine production by each NCS variant at 72 hours. Gray bar shows our original NCS from *Papaver somniferum*[2]. Error bars represent mean ± s.d. of three biological replicates.

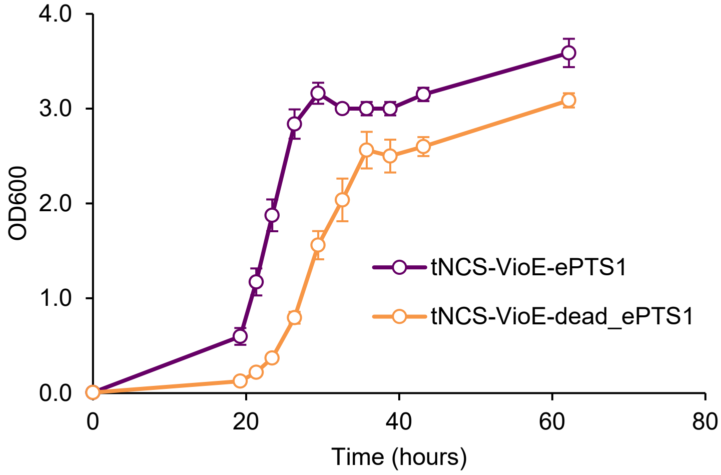


**Supplementary Figure 2. OD data from (*S*)-norcoclaurine shake flask fermentation experiment.** tNCS-VioE-ePTS1 and -dead_ePTS1 constructs were expressed using the strong pTDH3 promoter on a CEN6/ARS4 plasmid. Error bars represent mean ± s.d. of four biological replicates.

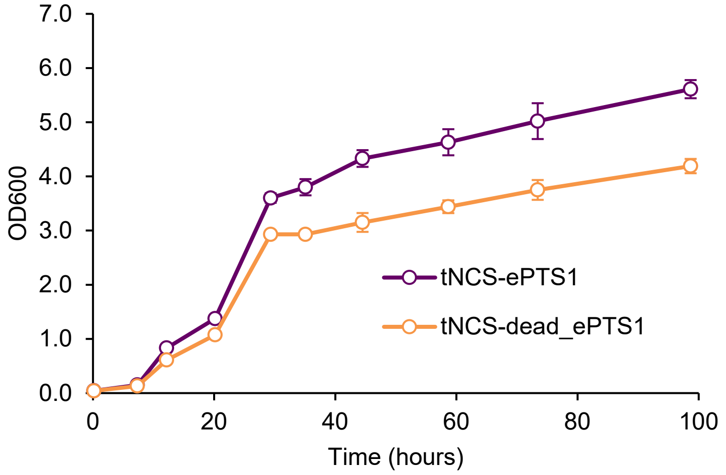


**Supplementary Figure 3. OD data from (*S*)-reticuline shake flask fermentation experiment.** tNCS-ePTS1 and -dead_ePTS1 constructs, along with 6OMT, CNMT, NMCH, and 4’OMT, were expressed on a CEN6/ARS4 plasmid. Error bars represent mean ± s.d. of four biological replicates.

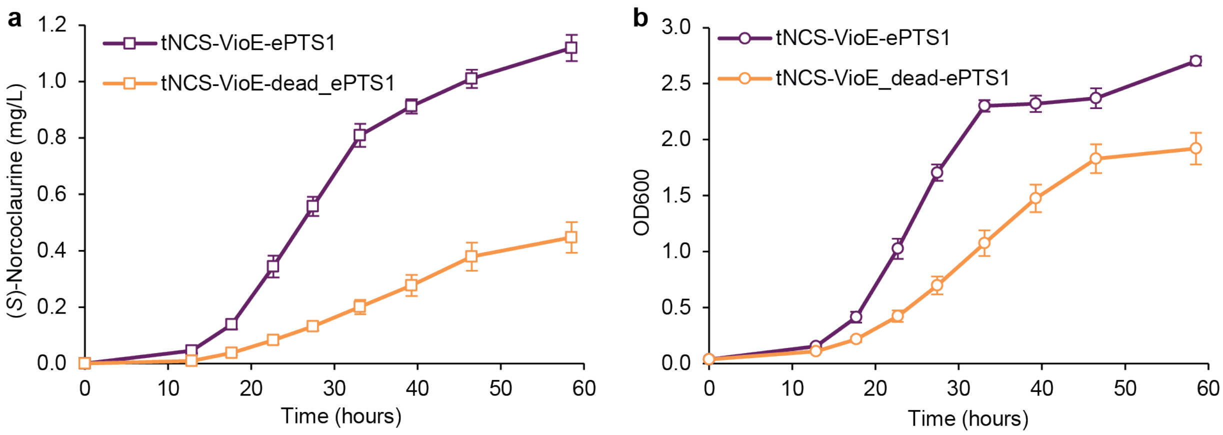


**Supplementary Figure 4. 2μ shake flask fermentation experiment.** (**a**) (*S*)-Norcoclaurine titer. (**b**) OD600. tNCS-VioE-ePTS1 and -dead_ePTS1 constructs were expressed using the strong pTDH3 promoter on a 2μ plasmid. Error bars represent mean ± s.d. of four biological replicates.

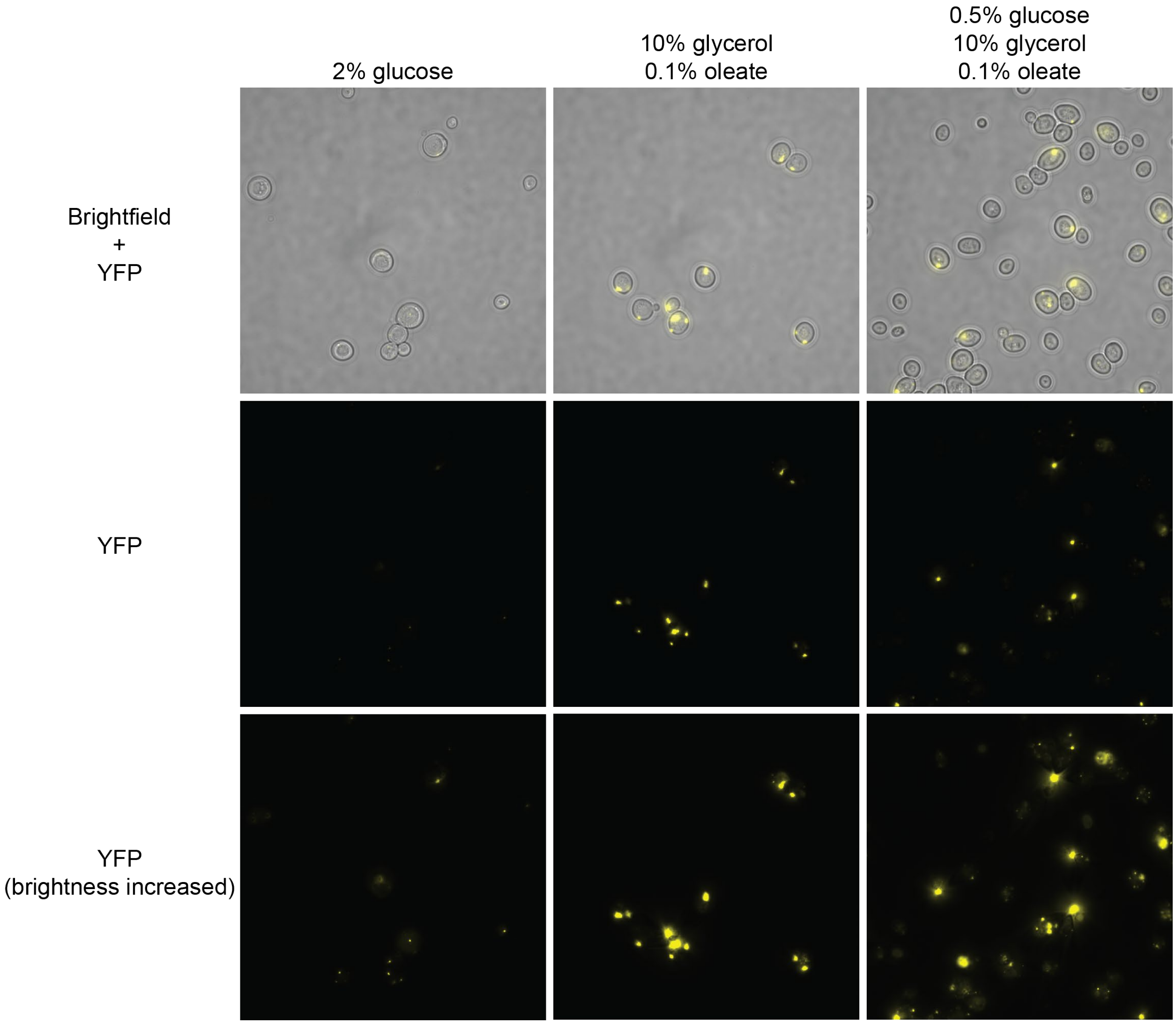


**Supplementary Figure 5. Oleate induction increases peroxisome size.** Peroxisomes were visualized using a strain expressing VioE-YFP-ePTS1 on a CEN6/ARS4 plasmid. In glucose, small peroxisomes were observed whereas larger peroxisomes were observed when cells were grown in oleate-containing media. Large peroxisomes are known to be induced when cells utilize fatty acids, including oleate[3,4]. All images were taken with identical exposure settings. After image acquisition, YFP channel brightness was increased during image processing to allow visualization of peroxisomes in glucose media (3rd row).

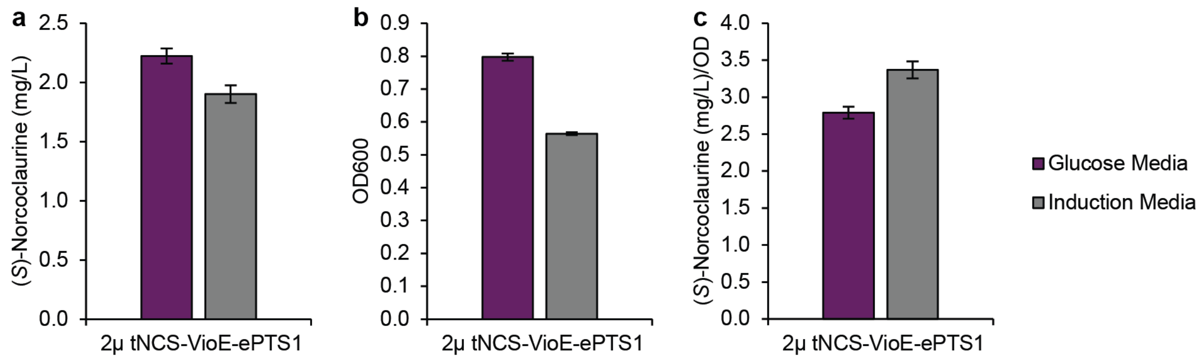


**Supplementary Figure 6. (*S*)-Norcoclaurine production using oleate induction media.** (**a**) (*S*)-Norcoclaurine titer. (**b**) OD600. (**c**) Titer normalized by OD. Glucose media is yeast synthetic complete media, minus leucine and uracil, plus 2% glucose. Induction media is our hybrid media: synthetic complete media, minus leucine and uracil, plus 0.5% glucose, 10% glycerol, 0.4% Tween80, and 0.1% oleate. Measurements were taken at 160 hours to allow sufficient time for growth and production. Error bars represent mean ± s.d. of four biological replicates.

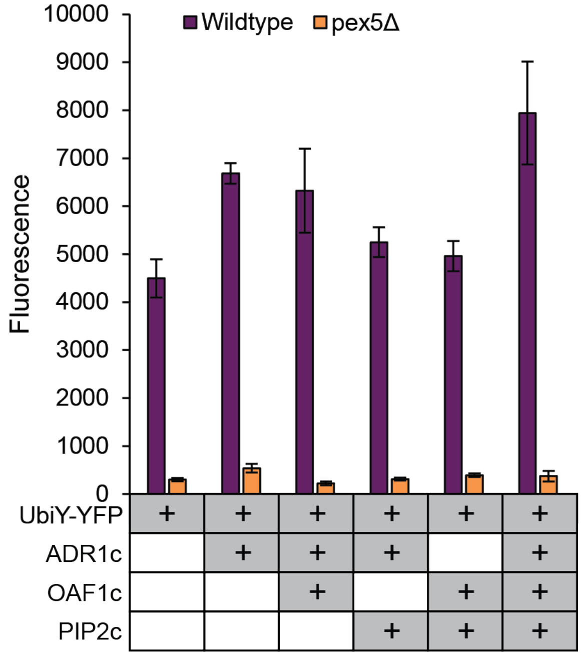


**Supplementary Figure 7. Expression of engineered transcription factors improves protection of peroxisomally-targeted UbiY-YFP-ePTS1.** Yellow fluorescent protein (YFP) was expressed with an N-terminal degradation tag (UbiY) in the presence or absence of constitutively-active transcription factors ADR1c, OAF1c, PIP2c. Wildtype strains import UbiY-YFP-ePTS1 into the peroxisome whereas pex5Δ strains are import-deficient due to knockout of the cytosolic receptor protein Pex5p. Samples were taken from a 96-well culture block for measurement at 48 hours. Error bars represent mean ± s.d. of five (UbiY-YFP only strains, first purple bar and first orange bar) or six (all others) biological replicates.

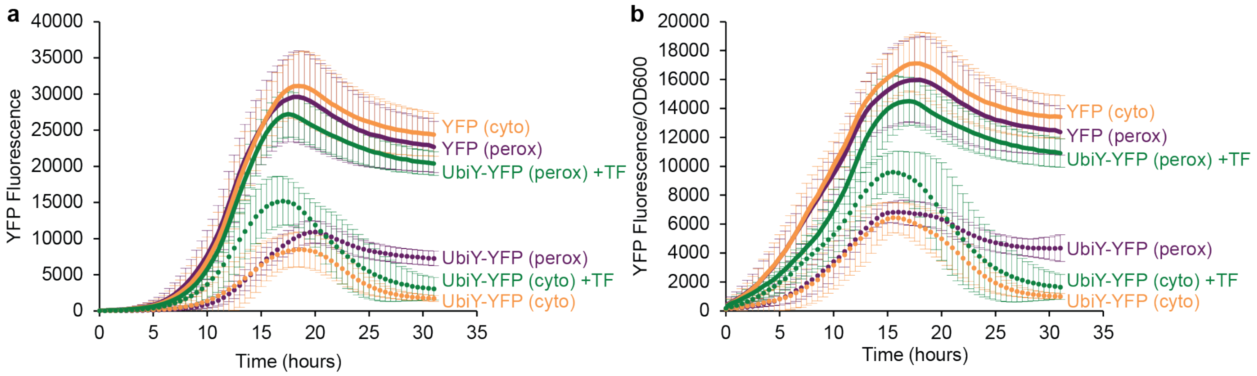


**Supplementary Figure 8. Dynamics of peroxisomal protection of UbiY-YFP-ePTS1 from degradation.** (**a**) Fluorescence. (**b**) Fluorescence normalized by OD600. Yellow fluorescent protein (YFP) was expressed with an N-terminal degradation signal (UbiY) in the presence or absence of constitutively-active transcription factors ADR1c/OAF1c/PIP2c (+TF). All strains contain the peroxisomal targeting signal ePTS1 fused to YFP. Constructs transformed into a wildtype background strain will target (UbiY-)YFP-ePTS1 protein to the peroxisome (perox). Cytosolic expression of (UbiY-)YFP-ePTS1 (cyto) is achieved by use of a pex5Δ background strain, which is import-deficient due to knockout of the cytosolic receptor protein Pex5p. Error bars represent mean ± s.d. of twelve biological replicates.

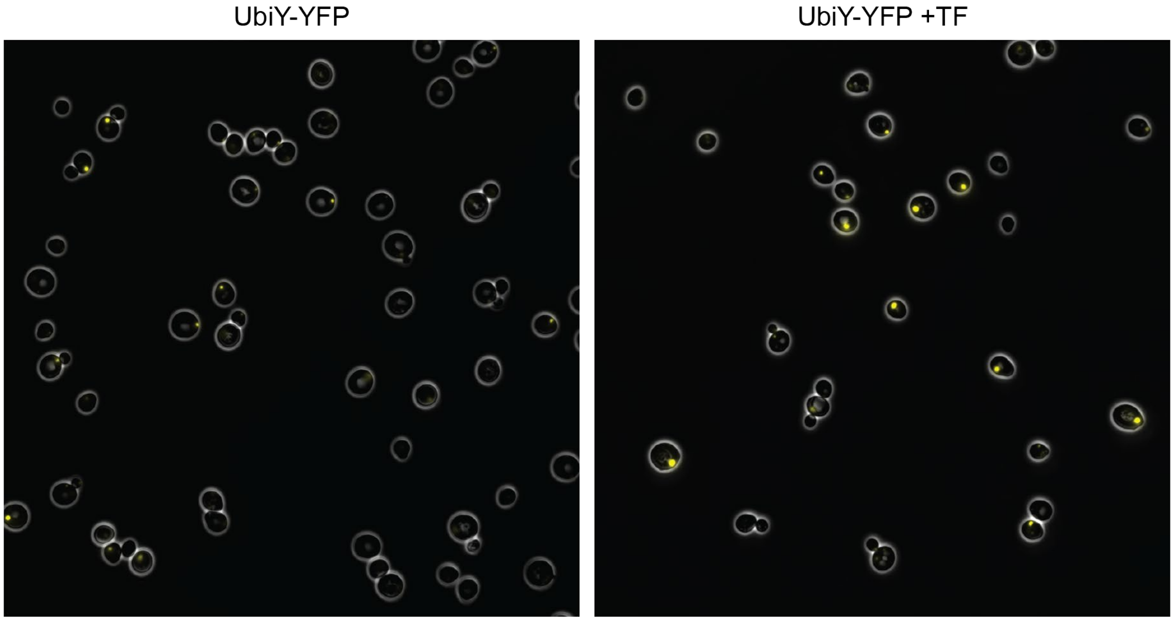


**Supplementary Figure 9. Expression of engineered transcription factors increases protection of peroxisomally-targeted UbiY-YFP-ePTS1 (zoomed-out version of Fig. 4c).** Fluorescence microscopy showing cells without (left) or with (right) expression of constitutively-active transcription factors ADR1c, OAF1c, PIP2c (+TF). Both strains contain YFP fused to the UbiY degradation signal on the N-terminus and the peroxisomal targeting signal ePTS1 on the C-terminus. YFP channel brightness was increased identically across both images to allow better visualization of peroxisomes from the UbiY-YFP strain.

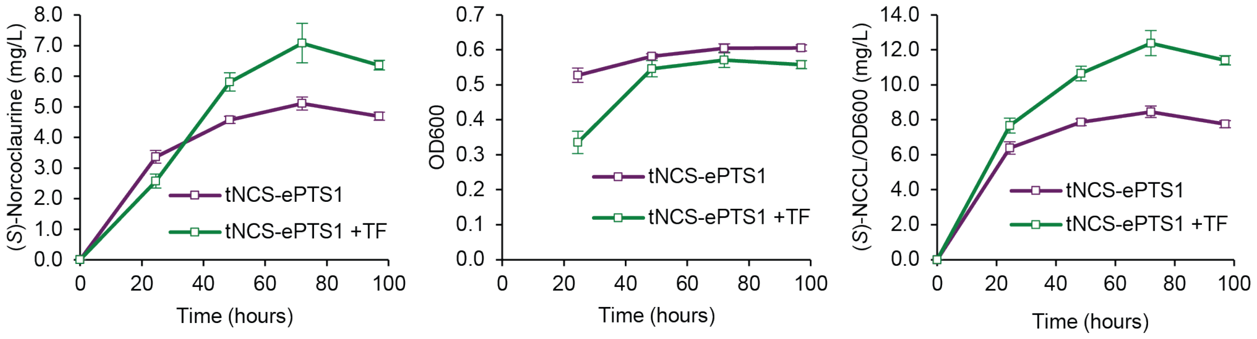


**Supplementary Figure 10. (*S*)-Norcoclaurine titer, OD600, and OD-normalized titer from transcription factor overexpression experiment.** All strains contain the upstream BIA pathway plus 2µ pTDH3-tNCS-ePTS1. The +TF strain also contains ADR1c, OAF1c, and PIP2c. Error bars represent mean ± s.d. of eight biological replicates.

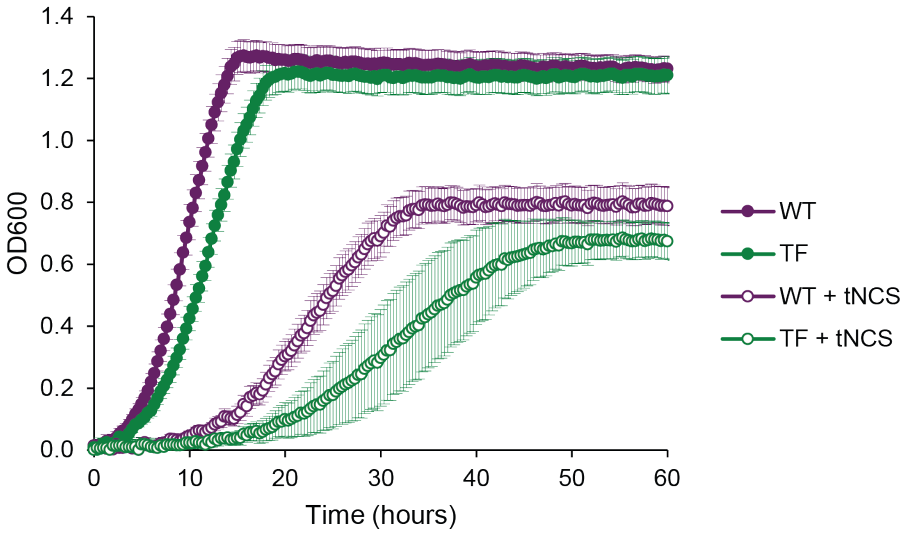


**Supplementary Figure 11. Expression of engineered transcription factors is linked to slower growth.** All strains contain the upstream BIA pathway. Strains without transcription factor overexpression (WT) contain a LEU2 marker. Strains with transcription factor overexpression (TF) contain ADR1c, OAF1c, PIP2c, and a LEU2 marker. Error bars represent mean ± s.d. of twelve biological replicates.

**Supplementary Table 1. Yeast Strains**

Plasmid sequences will uploaded with the final version of this manuscript.

| Strain # | Strain Name | Strain Parent | Plasmid Used | Description | Yeast Marker | Used in Figure |
|---|---|---|---|---|---|---|
| 1 | yWCD230 | BY4741 | pWCD1351 | his3Δ | | |
| 2 | yWCD231 | BY4741 pex5Δ::KanMX | pWCD1351 | his3Δ | KanMX | |
| 3 | yWCD745 | BY4741 | pWCD2249 | pTDH3-CYP76AD1_W13L_F309L-tTDH1-pCCW12-DODC-tADH1-pPGK1-ARO4_FBR-tPGK1 | URA3 | |
| 4 | yPSG633 | yWCD745 | pPSG958 | pTDH3-PsNCS-tENO2 | URA3, LEU2 | S1 |
| 5 | yPSG634 | yWCD745 | pPSG959 | pTDH3-PsNCSΔC-tENO2 | URA3, LEU2 | S1 |
| 6 | yPSG635 | yWCD745 | pPSG960 | pTDH3-TfNCS-tENO2 | URA3, LEU2 | S1 |
| 7 | yPSG636 | yWCD745 | pPSG961 | pTDH3-TfNCSΔN1-tENO2 | URA3, LEU2 | S1 |
| 8 | yPSG637 | yWCD745 | pPSG962 | pTDH3-TfNCSΔN2-tENO2 | URA3, LEU2 | S1 |
| 9 | yPSG638 | yWCD745 | pPSG963 | pTDH3-TfNCSΔN1ΔC-tENO2 | URA3, LEU2 | S1 |
| 10 | yPSG639 | yWCD745 | pPSG964 | pTDH3-TfNCSΔN2ΔC -tENO2 | URA3, LEU2 | S1 |
| 11 | yPSG627 | yWCD745 | pBSC009 | pTDH3-CjNCS-tENO2 | URA3, LEU2 | S1 |
| 12 | yPSG628 | yWCD745 | pPSG325 | pTDH3-CjNCSΔN1-tENO2 | URA3, LEU2 | S1 |
| 13 | yPSG629 | yWCD745 | pBSC011 | pTDH3-CjNCSΔN2-tENO2 | URA3, LEU2 | S1 |
| 14 | yPSG408 | yWCD231 | pPSG465 | pTDH3-CYP76AD5-tTDH1-pCCW12-DODC-tADH1-pPGK1-ARO4_FBR-tPGK1 | URA3 | |
| 15 | yPSG440 | yPSG408 | pPSG532 | CEN6/ARS4 | URA3, LEU2 | 1b, 1c, 1d |
| 16 | yPSG521 | yPSG408 | pPSG711 | pREV1-tNCS-ePTS1-tENO2-CEN6/ARS4 | URA3, LEU2 | 1b |
| 17 | yPSG520 | yPSG408 | pPSG710 | pRNR2-tNCS-ePTS1-tENO2-CEN6/ARS4 | URA3, LEU2 | 1b |
| 18 | yPSG519 | yPSG408 | pPSG709 | pRPL18B-tNCS-ePTS1-tENO2-CEN6/ARS4 | URA3, LEU2 | 1b, 1c |
| 19 | yPSG518 | yPSG408 | pPSG708 | pTEF1-tNCS-ePTS1-tENO2-CEN6/ARS4 | URA3, LEU2 | 1b |
| 21 | yPSG438 | yPSG408 | pPSG569 | pTDH3-tNCS-ePTS1-tENO2-CEN6/ARS4 | URA3, LEU2 | 1b, 1c, 1d |
| 22 | yPSG723 | yWCD231 | pPSG569 | pTDH3-tNCS-ePTS1-tENO2-CEN6/ARS4 | LEU2 | 1d |
| 23 | ySJD004 | yWCD230 | pSJD001 | YBR197CΔ::pPAB1-Pex22(1-36)-mRuby2-tSSA1 | | |

| | | | | | | |
|---|---|---|---|---|---|---|
| 24 | yPSG751 | ySJD004 | pPSG1066 | pTDH3-tNCS-Venus-ePTS1-tENO2-CEN6/ARS4 | LEU2 | 2b |
| 25 | yPSG306 | yWCD230 | pPSG465 | pTDH3-CYP76AD5-tTDH1-pCCW12-DODC-tADH1-pPGK1-ARO4_FBR-tPGK1 | URA3 | |
| 26 | yPSG575 | yPSG306 | pPSG804, pPSG820 | YDR514CΔ::pTDH3-VioA-tENO1-pTEF1-VioB-tPGK1 | URA3 | |
| 27 | yJAS816 | yPSG575 | pJAS1643 | pTDH3-tNCS-VioE-ePTS1-tENO2-CEN6/ARS4 | URA3, LEU2 | 2c, 2d, 2e, S2 |
| 28 | yJAS817 | yPSG575 | pJAS1644 | pTDH3-tNCS-VioE-dead_ePTS1-tENO2-CEN6/ARS4 | URA3, LEU2 | 2c, 2d, 2e, S2 |
| 29 | yJAS832 | yPSG575 | pJAS1668 | pTDH3-VioE-ePTS1-tADH1-CEN6/ARS4 | URA3, LEU2 | 2d |
| 30 | yJAS833 | yPSG575 | pJAS1669 | pTDH3-VioE-dead_ePTS1-tADH1-CEN6/ARS4 | URA3, LEU2 | 2d |
| 31 | yPSG718 | yPSG306 | pPSG1059 | pTDH3-tNCS-ePTS1-tENO2-pTEF1-Ec_CYP80B1-tTDH1-pCCW12-Ps_6OMT-tPGK1-pPGK1-Ps_4'OMT2-tENO1-pTEF2-Ps_CNMT-tSSA1-CEN6/ARS4 | URA3, LEU2 | 2g, S3 |
| 32 | yPSG719 | yPSG306 | pPSG1060 | pTDH3-tNCS-dead_ePTS1-tENO2-pTEF1-Ec_CYP80B1-tTDH1-pCCW12-Ps_6OMT-tPGK1-pPGK1-Ps_4'OMT2-tENO1-pTEF2-Ps_CNMT-tSSA1-CEN6/ARS4 | URA3, LEU2 | 2g, S3 |
| 33 | yPSG752 | ySJD004 | pPSG1067 | pTDH3-tNCS-Venus-ePTS1-tENO2-2μ | LEU2 | 3a |
| 34 | yJAS971 | yPSG575 | pJAS1766 | pTDH3-VioE-ePTS1-tENO2-2μ | URA3, LEU2 | 3c |
| 35 | yJAS972 | yPSG575 | pJAS1767 | pTDH3-VioE-dead_ePTS1-tENO2-2μ | URA3, LEU2 | 3c |
| 36 | yJAS973 | yPSG575 | pJAS1769 | pTDH3-tNCS-VioE-ePTS1-tENO2-2μ | URA3, LEU2 | 3b, 3c, 3d, S4, S6 |
| 37 | yJAS974 | yPSG575 | pJAS1770 | pTDH3-tNCS-VioE-dead_ePTS1-tENO2-2μ | URA3, LEU2 | 3b, 3c, 3d, S4 |
| 38 | yPSG335 | yWCD230 | pJAS1052 | pTDH3-VioA-tENO1-pTEF1-VioB-tPGK1 | URA3 | |
| 39 | yPSG666 | yPSG335 | pPSG1010 | pTEF1-VioE-Venus-ePTS1-tENO2-CEN6/ARS4 | URA3, LEU2 | S5 |
| 40 | yJJB006 | yWCD230 | pJJB111 | pTDH3-Venus-ePTS1-tTDH1 | URA3 | |
| 41 | yJJB012 | yWCD231 | pJJB111 | pTDH3-Venus-ePTS1-tTDH1 | URA3 | |
| 42 | yJJB013 | yWCD230 | pJJB207 | pTDH3-UbiY-Venus-ePTS1-tTDH1 | URA3 | |
| 43 | yJJB014 | yWCD231 | pJJB207 | pTDH3-UbiY-Venus-ePTS1-tTDH1 | URA3 | |
| 44 | yPSG777 | yJJB006 | pPSG372 | LEU2 marker | URA3, LEU2 | 4b, S8 |
| 45 | yPSG778 | yJJB012 | pPSG372 | LEU2 marker | URA3, LEU2 | 4b, S8 |

| | | | | | | |
|---|---|---|---|---|---|---|
| 46 | yJJB150 | yJJB013 | pPSG372 | LEU2 marker | URA3,<br>LEU2 | 4b, 4c,<br>S7,<br>S8, S9 |
| 47 | yJJB151 | yJJB014 | pPSG372 | LEU2 marker | URA3,<br>LEU2 | 4b,<br>S7, S8 |
| 48 | yJJB154 | yJJB013 | pJJB306 | pRPL18B-ADR1c-tPGK1 | URA3,<br>LEU2 | S7 |
| 49 | yJJB157 | yJJB014 | pJJB306 | pRPL18B-ADR1c-tPGK1 | URA3,<br>LEU2 | S7 |
| 50 | yJJB064 | yJJB013 | pJJB175 | pRPL18B-ADR1c-tPGK1-<br>pRPL18B-OAF1c-tTDH2 | URA3,<br>LEU2 | S7 |
| 51 | yJJB076 | yJJB014 | pJJB175 | pRPL18B-ADR1c-tPGK1-<br>pRPL18B-OAF1c-tTDH2 | URA3,<br>LEU2 | S7 |
| 52 | yJJB061 | yJJB013 | pJJB172 | pRPL18B-ADR1c-tPGK1-<br>pRPL18B-PIP2c-tTEF1 | URA3,<br>LEU2 | S7 |
| 53 | yJJB073 | yJJB014 | pJJB172 | pRPL18B-ADR1c-tPGK1-<br>pRPL18B-PIP2c-tTEF1 | URA3,<br>LEU2 | S7 |
| 54 | yJJB069 | yJJB013 | pJJB180 | pRPL18B-PIP2c-tTEF1-<br>pRPL18B-OAF1c-tTDH2 | URA3,<br>LEU2 | S7 |
| 55 | yJJB081 | yJJB014 | pJJB180 | pRPL18B-PIP2c-tTEF1-<br>pRPL18B-OAF1c-tTDH2 | URA3,<br>LEU2 | S7 |
| 56 | yJJB059 | yJJB013 | pJJB170 | pRPL18B-ADR1c-tPGK1-<br>pRPL18B-PIP2c-tTEF1-<br>pRPL18B-OAF1c-tTDH2 | URA3,<br>LEU2 | 4b, 4c,<br>S7,<br>S8, S9 |
| 57 | yJJB071 | yJJB014 | pJJB170 | pRPL18B-ADR1c-tPGK1-<br>pRPL18B-PIP2c-tTEF1-<br>pRPL18B-OAF1c-tTDH2 | URA3,<br>LEU2 | 4b,<br>S7, S8 |
| 58 | yPSG726 | yPSG306 | pPSG372 | LEU2 marker | URA3,<br>LEU2 | S11 |
| 59 | yPSG727 | yPSG306 | pJJB170 | pRPL18B-ADR1c-tPGK1-<br>pRPL18B-PIP2c-tTEF1-<br>pRPL18B-OAF1c-tTDH2 | URA3,<br>LEU2 | S11 |
| 60 | yPSG738 | yPSG726 | pPSG830 | pTDH3-tNCS-ePTS1-tENO2-<br>2μ | URA3,<br>LEU2,<br>HIS3 | 4d,<br>S10,<br>S11 |
| 61 | yPSG739 | yPSG727 | pPSG830 | pTDH3-tNCS-ePTS1-tENO2-<br>2μ | URA3,<br>LEU2,<br>HIS3 | 4d,<br>S10,<br>S11 |
